# PyHIST: A Histological Image Segmentation Tool

**DOI:** 10.1101/2020.05.07.082461

**Authors:** Manuel Muñoz-Aguirre, Vasilis F. Ntasis, Roderic Guigó

## Abstract

The development of increasingly sophisticated methods to acquire high resolution images has led to the generation of large collections of biomedical imaging data, including images of tissues and organs. Many of the current machine learning methods that aim to extract biological knowledge from histopathological images require several data preprocessing stages, creating an overhead before the proper analysis. Here we present PyHIST (https://github.com/manuel-munoz-aguirre/PyHIST), an easy-to-use, open source whole slide histological image tissue segmentation and preprocessing tool aimed at data preparation for machine learning applications.

## Main text

In histopathology, Whole Slide Images (WSI) are high resolution images of tissue sections obtained by scanning conventional glass slides. Histopathological images are routinely used in the diagnosis of many diseases, notably cancer. The increasing automation of WSI acquisition has led to the development of computational methods to process the images and help clinicians and pathologists in diagnosis and disease classification. As an increasing number of larger WSI datasets became available, methods have been developed for a wide array of tasks, such as the classification of breast cancer metastases, Gleason scoring for prostate cancer, tumor segmentation, nuclei detection and segmentation, bladder cancer diagnosis, mutated gene prediction based on pathology images, among others ^1–6^. Besides of being important diagnostic tools, histopathological images capture endophenotypes (of organs and tissues) that, when correlated with molecular and cellular data on the one hand, and higher order phenotypic traits on the other, can provide crucial information on the biological pathways that mediate between the sequence of the genome and the biological traits of the organisms (including diseases).

Because of the complexity of the information typically contained in WSIs, Machine Learning (ML) methods that can infer, without prior assumptions, the relevant features that they encode are becoming the preferred analytical tools. These features may be clinically relevant but challenging to spot even for expert pathologists, and thus, ML methods can prove valuable in healthcare decision-making ^7^.

In most ML tasks, data preprocessing remains a fundamental step. Indeed, in the domain of histological images, there are a number of issues when preprocessing the data before an analysis: due to the large dimensions of WSIs, many deep learning applications have to break them down into smaller-sized square pieces called tiles ^8^. Furthermore, a significant fraction of the area in a WSI is often background, which does not contain useful information and can cause computing overhead in downstream analyses. To circumvent this, some applications apply a series of image transformations to identify the foreground from the background (see, for example, ^9^), and perform relevant operations only over regions with tissue content. However, this process is not standardized, and customized scripts have to be frequently developed to deal with data preparation stages. This is cumbersome and may introduce dataset specific-biases, which can prevent integration across multiple datasets.

To systematize the WSI preprocessing procedure for ML applications, we developed PyHIST, a pipeline to segment the regions of a histological image into tiles with relevant tissue content (foreground) with little human intervention.

PyHIST is a Python tool based on OpenSlide ^10^, a library to read high-resolution histological images in a memory-efficient way. PyHIST’s input is a WSI encoded in SVS format (Fig. 1a), and the main output is a series of image tiles retrieved from regions with tissue content (Fig. 1e).

**Figure 1:**
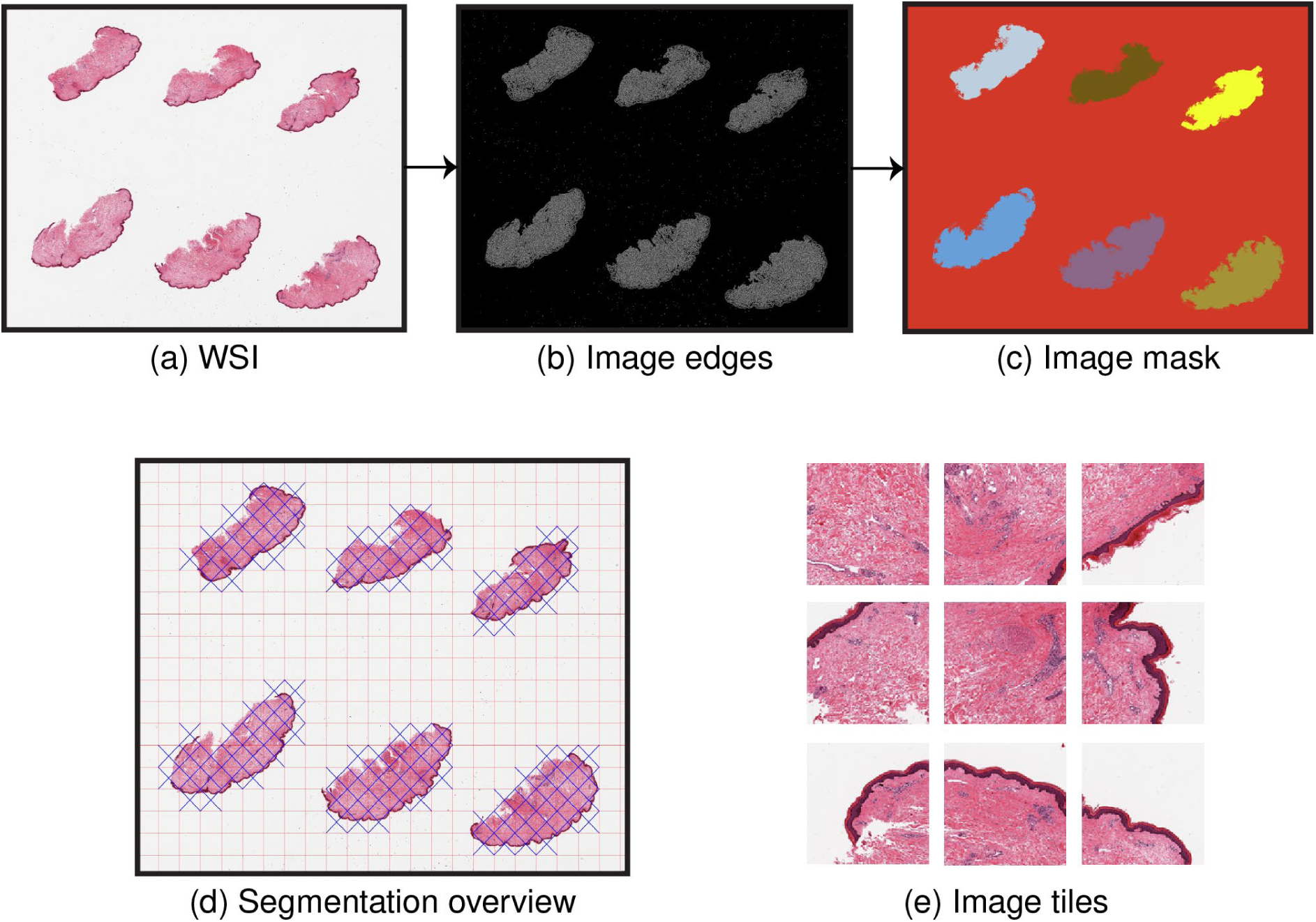
PyHIST pipeline. **(a)** The input to the pipeline is a Whole Slide Image (WSI). Within PyHIST, the user can decide to scale down the image to perform the segmentation and tile extraction at lower resolutions. The WSI shown is of a skin tissue sample (GTEX-1117F-0126) from the Genotype-Tissue Expression (GTEx) project ^12^ **(b)** An alternate version of the input image is generated, where the tissue edges are highlighted using a Canny edge detector. A graph segmentation algorithm is employed over this image, in order to generate the mask shown in **(c)**. PyHIST extracts tiles of specific dimensions from the masked regions, and provides an overview image to inspect the output of the segmentation and masking procedure, as shown in **(d)**, where the red lines indicate the grid generated by tiling the image at user-specified tile dimensions, while the blue crosses indicate the selected tiles meeting a certain user-specified threshold of tissue content with respect to the total area of the tile. In **(e)**, examples of selected tiles are shown.

The PyHIST pipeline involves three main steps: first, tissue edges inside the WSI are identified using a Canny edge detector (Fig. 1b), generating an alternate version of the image with diminished noise and an enhanced distinction between the background and the tissue foreground. Second, these edges are processed by a graph-based segmentation algorithm ^11^, which is used here to identify tissue content. In short, this step evaluates the boundaries between different regions of an image as defined by the edges; different parts of the image are represented as connected components of a graph and the “within” and “in-between” variations of neighbouring components are assessed in order to decide if the examined image regions should be merged or not into a single component. From this, a mask is obtained in which the background and the different tissue slices are separated and marked as distinct objects using different colors (Fig. 1c). Finally, the selected regions are divided into tiles at a user-specified size. Optionally, it is possible to only select tiles that contain a proportion of tissue above a given threshold with respect to the total area of the tile.

Of note, tile generation can be performed at the native resolution of the WSI, but downsampling factors can also be specified to generate tiles at lower resolutions. Additionally, edge detection and mask generation can also be performed on downsampled versions of WSI - reducing segmentation runtimes (Fig. S1, Methods). A segmentation overview image is generated at the end of the segmentation procedure for the user to visually inspect the selected tiles (Fig. 1d). With the set of parameters available in PyHIST (Supplementary text), the user can specify regions to ignore when performing the masking and segmentation (Fig. S2), and have a fine grained control over specific use-cases. PyHIST also has a random tile sampling mode for those applications that do not necessarily need to distinguish the background from the foreground. In this mode, tiles at a user-specified size and resolution will be extracted from random starting positions in the WSI.

To demonstrate how PyHIST can be used to preprocess WSIs for usage in a ML application, we generated a use case example with the goal of building a classifier at the tile-level that allows us to determine the tissue of origin based on the histological patterns encoded in these tiles. To this end, we first retrieved a total of 30 publicly available WSIs, 5 from each of the following human tissues hosted in The Cancer Genome Atlas (TCGA) ^13^: Brain, Breast, Colon, Kidney, Liver, and Skin. Second, these WSIs were preprocessed with PyHIST, generating a total of 4720 tiles with dimensions 512×512. These tiles were then partitioned into training and test sets, and we then fit a deep learning model over these tiles, achieving a classification accuracy of 94% (Fig. 2a, Table S1). We also inspected the feature vectors generated by the deep learning model: for each tile, we retrieved the features corresponding to the last layer of the network, and performed dimensionality reduction (tSNE) over the stacked matrix of these vectors. From here, we infer that the learned features clearly recapitulate tissue morphology since tile clusters corresponding to each tissue are formed (Figs. 2b, S3).

**Figure 2:**
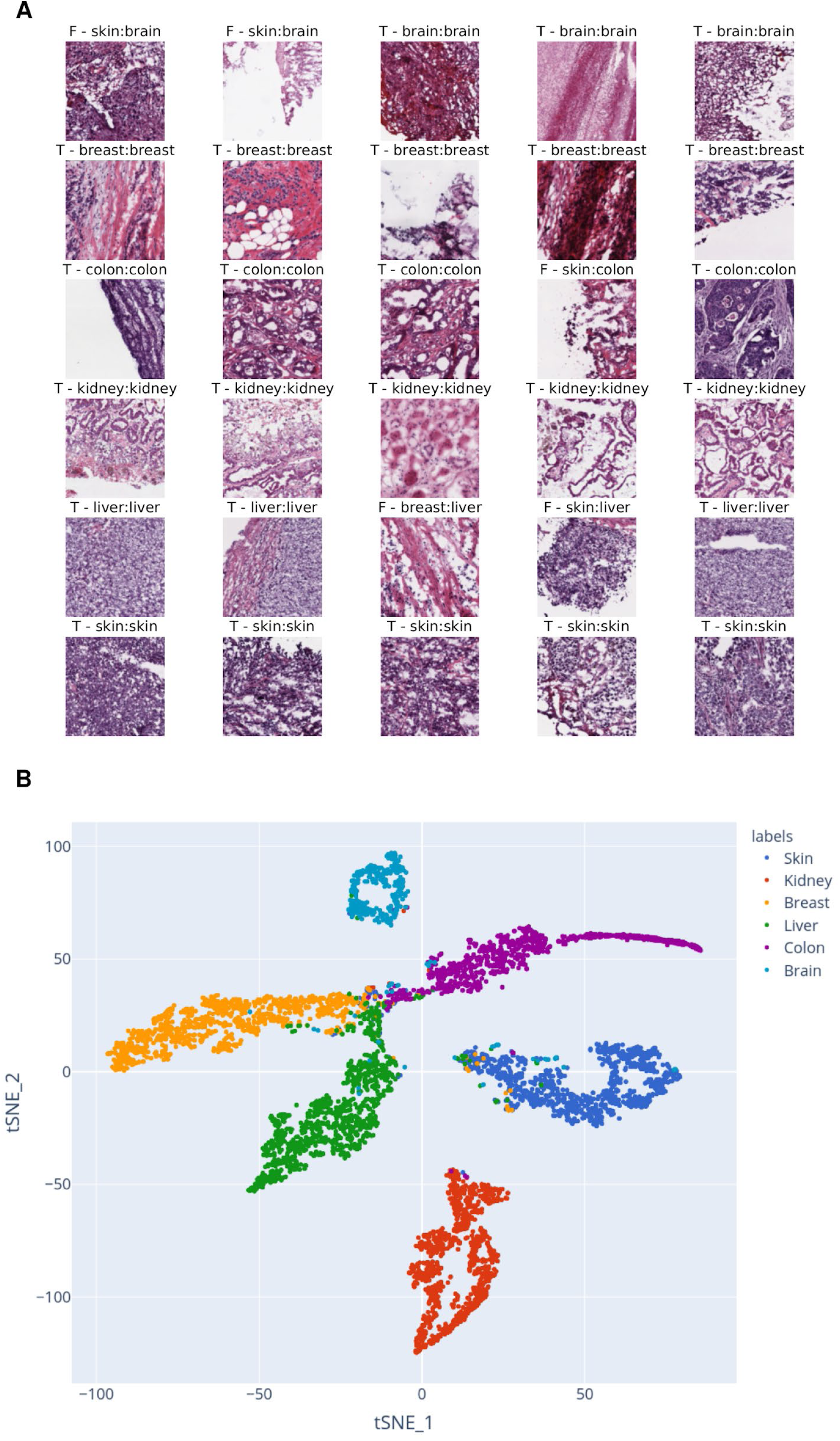
TCGA use case. **(a)** Random examples of tile predictions from TCGA tissue tiles. 30 random tiles are shown, 5 tiles per tissue in each row. Above each tile, “T” indicates that the prediction was correct and “F” incorrect, followed by the predicted tissue label, and the ground truth tissue label for that tile. **(b)** Dimensionality reduction of TCGA tiles. tSNE performed with the feature vectors (activations) of each tile that were derived from the deep learning classifier model. Each dot corresponds to an image tile.

With this use case, we have shown how to quickly prepare WSI data using PyHIST without manually preprocessing the images, reducing the overhead to start performing downstream analyses. The example use case described here is documented and fully available at https://pyhist.readthedocs.io/en/latest/testcase/, and divided into three Jupyter notebooks: 1) Data preprocessing with PyHIST, 2) Constructing a deep learning tissue classifier, and 3) Feature exploration.

PyHIST is a generic tool to segment histological images automatically: it allows for easy and rapid WSI cleaning and preprocessing with minimal effort to generate image tiles geared towards usage in ML analyses. The tool is highly customizable, enabling the user to tune the segmentation process in order to suit the needs of any particular application that relies on histological image tiles. PyHIST and all of its dependencies have been packaged in a Docker image, ensuring portability across different systems. PyHIST can also be used locally on a regular computing environment with minimal requirements. Finally, PyHIST is open source software: all the code and reproducible notebooks for the example use case are available in GitHub and will continue to be improved on the basis of user feedback.

## Methods

### Overview of PyHIST

PyHIST is a high-resolution histological image segmentation pipeline. It takes as input a Whole Slide Image (WSI) encoded in SVS format (Fig. 1a) and produces a series of image tiles (Fig. 1e), with dimensions specified by the user. These tiles are usually extracted from the regions of the WSI that have tissue content, after discarding the background, although it is also possible to extract all the tiles from the WSI. Tile extraction is performed as if overlaying a grid with tiles of a fixed size over the original image (Fig. 1d), and then keeping the relevant tiles. It is also possible to sample tiles from random starting positions in the WSI. In both cases, tile extraction can be performed either at the original resolution of the image, or at a downsampled version.

### WSI segmentation

In order to identify the sections of the WSI that contain tissue, PyHIST first uses a Canny edge detector ^14^ to generate a version of the image that highlights edges within tissue fragments. This image is then used in combination with a graph segmentation algorithm ^11^ to select connected components in the image, obtaining a clean mask that corresponds to tissue segments (Fig. 1c). The mask will be then divided into tiles. Each tile will be inspected to determine if it’s composed by at least a certain percentage of tissue content with respect to the total area of the tile (see argument *--content-threshold*, Supplementary text). If the tile is kept, then the corresponding area of the original WSI will be retrieved. Tiles can be saved in JPG or PNG formats.

Although it is possible to perform the segmentation at the highest resolution encoded in the WSI, by default PyHIST will output tiles from a downsampled version (i.e. a lower resolution version of the image, scaled by a factor). The user can specify a downsampling factor (powers of 2) to perform tile extraction at a lower resolution (see argument *--output-downsample*, Supplementary text). Of note, incomplete tiles can be generated towards the right and lower borders of the image due to the selected tile size not being a multiple of the WSI size. In this case, the size of these tiles will be as large as possible, without exceeding the specified tile dimension. When the process is finished, a tab-delimited text file will be also produced indicating tile coordinates and size of each tile, as well as an indicator column stating if the tile passed the content threshold to be considered foreground.

Several options are provided to tune PyHIST for specific segmentation use-cases. Usually, tissue segments are placed towards the center of the slide during the imaging process. However, in some cases, tissue can be located in the borders of the slide, and depending on the application, the user would like to keep or remove these regions. PyHIST provides parameters to deal with these types of cases (Supplementary text).

It is also possible to extract a given number of tiles randomly from the WSI at any given resolution. In this case, since the tiles are sampled from any position, there is no need to perform segmentation.

### Intermediate scaling

To speed up the segmentation process, the input image (Fig. S1a) can be downscaled at different resolutions, at different steps of the pipeline, not only initially. If the mask is downscaled (Fig. S1b), the edge generation and segmentation process will be performed at a lower resolution, reducing the time needed to complete the process. The output resolution used to generate the tiles (Fig. S1c) is independent of all other choices: for example, a user can decide to generate the mask at 16x, but output tiles at native (1x) or any other resolution. If the mask is closer to the native resolution, a more precise segmentation is obtained, however, for most applications, such level of detail is not necessary. The same is applicable for the segmentation overview image (Fig. S1d), which can be generated at any resolution independently of the other choices. All these arguments are available in the “Downsampling” section of the Supplementary text.

### Test mode

A test mode (argument *--test-mode*) is available in PyHIST to inspect how a mask will look with a given combination of segmentation parameters, as well as to verify the tiling grid that will be generated at the selected tile dimensions (Fig. S2). This is to aid the user in inspecting the output before producing the individual tile files. When the test mode is invoked, no tile selection will be performed, as it only serves to assess how the tiles will be generated.

### Execution times

We benchmarked PyHIST’s execution time both for random tile sampling and segmentation. Using the WSI in Fig. S1 with native dimensions 47807×38653, we performed random sampling of a varying number of tiles at different downsampling factors (Fig. S4a), observing a linear trend in runtime with respect to the number of tiles. Efficiency will be determined by the resolution levels natively encoded in the WSI generation process: for this WSI, 1x, 4x and 16x are available. If the requested downsampling factors are available in the image layers of the WSI file, segmentation will be faster. On the other hand, if the requested downsampling factor is not encoded in the WSI (like 8x in this example), sampling will run for a longer time since resizing operations need to be performed. We also examined the runtime of performing random sampling of 1000 tiles at a fixed tile size, and at different downsampling levels (Fig. S4b). As expected, runtime increases with tile size. Similarly to the previous benchmark, here we also observe the effect that the native encodings in the WSI have on the runtime speed: since 8x is not encoded in the WSI, resizing operations need to be performed, leading to longer runtimes when compared to 1x and 4x.

To benchmark the segmentation process, we use 50 different WSIs of stomach tissue (with different dimensions) from the Genotype-Tissue Expression (GTEx) project ^12^ and perform segmentation at different downsampling factors at a fixed tile size of 256×256. Runtime variability within a single downsampling factor will be determined by slide size; and segmentation runtime decreases almost linearly with respect to resolution (Fig. S4c), while the runtime will increase linearly with the dimension of the slide (Fig. S4d).

### TCGA tissue classification use case

We downsample the original WSIs by a factor of 4x, and require that a tile is composed of at least 40% of tissue content in order to consider it for further analysis. From the 30 WSIs, a total of 4720 tiles with dimensions 512×512 were produced. We performed data augmentation by applying a set of transformations to the image tiles (rotations, resizing, crops, and flips). We then use a ResNet152 ^15^ deep learning architecture pretrained on ImageNet ^16^ to classify the tissue of origin for each tile (Fig. 2a). Transfer learning is performed by changing the last fully connected layer of the model, freezing the rest of the network, and training only the last layer.

## Acknowledgments

We acknowledge Kaiser Co and Valentin Wucher for testing PyHIST, and the colleagues at the lab for useful feedback. This work was supported by FPU15/03635, Ministerio de Educación, Cultura y Deporte to M.M-A. All authors acknowledge the support of the Spanish Ministry of Science, Innovation and Universities to the EMBL partnership, the Centro de Excelencia Severo Ochoa, and the CERCA Programme / Generalitat de Catalunya.

## Author contributions

M.M.-A. and R.G. conceived the idea of PyHIST. V.F.N. implemented the core features of PyHIST. M.M.-A. implemented usability features, performance upgrades and packaged the Docker image for PyHIST. V.F.N. and M.M.-A. conceived the TCGA example use case and implemented reproducible notebooks. R.G. provided conceptual advice. All authors participated in writing the manuscript.

## Competing interests

The authors declare no competing interests.

## Code availability

PyHIST is available at https://github.com/manuel-munoz-aguirre/PyHIST and released under a GPL license. Updated documentation and a tutorial can be found at https://pyhist.readthedocs.io/

## Data availability

The TCGA WSIs in the Use Case were downloaded from the Genomic Data Commons (GDC) repository (https://gdc.cancer.gov/) using the GDC Data-transfer tool (https://gdc.cancer.gov/access-data/gdc-data-transfer-tool).

## Notes

https://github.com/manuel-munoz-aguirre/PyHIST

### Competing Interest Statement

The authors have declared no competing interest.

