## Supplementary material for "PyHIST: A Histological Image Segmentation Tool"

### Parameter description

The parameters in PyHIST are divided into five main functional blocks:

**Positional arguments:**

| | |
|---|---|
| input_image | The whole slide image input file |

**Optional arguments:**

| | |
|---|---|
| -h, --help | show this help message and exit |

**Execution:** related to how the pipeline will execute and extract the patches.

| | |
|---|---|
| --patch-size PATCH_SIZE | Integer indicating the size of the produced patches. A value of P will produce patches of size P x P. (default: 512) |
| --format {png,jpg} | Format to save the patches. (default: png) |
| --verbose | Print status messages at each step of the pipeline (both for segmentation and sampling (default: False) |
| --test-mode | Trigger test mode for image mask and tile debugging. (default: False) |

**Sampling:** related to patch extraction from random starting positions in the WSI, instead of a fixed grid.

| | |
|---|---|
| --sampling | Random patch sampling mode. (default: False) |
| --npatches NPATCHES | Number of patches to extract in random sampling mode. (default: 100) |

**Output:** related to patch saving, as well as other intermediate images generated by PyHIST that can be helpful for debugging and output inspection.

| | |
|---|---|
| --output OUTPUT | Output directory (default: output/) |
| --save-tilecrossed-image | Produce a thumbnail of the original image, in which the selected tiles are marked with a cross. (default: False) |
| --save-edges | Keep the image produced by the Canny edge detector. (default: False) |
| --save-mask | Keep the mask with tissue segments. (default: False) |
| --save-patches | Save all the produced tiles of the full resolution image. (default: False) |
| --exclude-blank | If enabled, background tiles will not be saved. (default: False) |
| --save-nonsquare | By default, only square tiles are saved, discarding the regions towards the edges of the WSI that do not fit a complete tile. If this flag is enabled, these |

```
                        non-square tiles will be saved as well. (default:
                        False)
```

**Downsampling:** related to downsampling factors used to scale down the WSI resolution at different points of the pipeline.

```
  --output-downsample OUTPUT_DOWNSAMPLE
                        Downsampling factor for the output image. Must be a
                        power of 2. (default: 16)
  --mask-downsample MASK_DOWNSAMPLE
                        Downsampling factor to calculate the image mask. A
                        higher number will make the mask computer faster at
                        the expense of segmentation quality. Must be a power
                        of 2. (default: 16)
  --tilecross-downsample TILECROSS_DOWNSAMPLE
                        Downsampling factor to generate the tilecrossed
                        overview image. Must be a power of 2. (default: 16)
  --test-downsample TEST_DOWNSAMPLE
                        Downsampling factor to calculate the test image. Must
                        be a power of 2. (default: 16)
```

**Segmentation:** parameters to tune how the segmentation will be performed.

```
  --borders
{1001,1111,1000,0010,1110,0001,0011,0101,0111,0110,1010,1100,1101,0100,0000,1011}
                        A four digit string. Each digit represents a border of
                        the image in the following order: left, bottom, right,
                        top. If the digit is 1, then the corresponding border
                        will be taken into account to define background. For
                        instance, with 1010 the algorithm will look at the
                        left and right borders of the segmented image, in a
                        window of width defined by the --percentage-bc
                        argument, and every segment identified will be set as
                        background. This argument is mutually exclusive with
                        --corners. If --borders is set to be different from
                        0000, then --corners must be 0000. Default value is
                        1111. (default: 1111)
  --corners
{1001,1111,1000,0010,1110,0001,0011,0101,0111,0110,1010,1100,1101,0100,0000,1011}
                        A four digit string. Each digit represents a corner of
                        the image in the following order: top left, bottom
                        left, bottom right, top right. If the digit is equal
                        to 1, then the corresponding corner will be taken into
                        account to define background. For instance, with 0101,
                        the bottom left and top right corners of the segmented
                        image will be considered, with a square window of size
```

given by the --percentage-bc argument, and every segment identified will be set as background. This argument is mutually exclusive with --borders. If --corners is set to be different from 0000, then --borders must be 0000. Default value is 0000. (default: 0000)

--k-const K_CONST

Parameter required by the segmentation algorithm. Value for the threshold function. The threshold function controls the degree to which the difference between two segments must be greater than their internal differences in order for them not to be merged. Lower values result in finer segmentation. Larger images require higher values. Default value is 10000. (default: 10000)

--minimum_segmentsize MINIMUM_SEGMENTSIZE

Parameter required by the segmentation algorithm. Minimum segment size enforced by post-processing. Larger images require higher values. Default value is 10000. (default: 10000)

--percentage-bc PERCENTAGE_BC

Integer [0-100] indicating the percentage of the image (width and height) that will be considered as border/corner in order to define the background. (default: 5)

--sigma SIGMA

Parameter required by the segmentation algorithm. Used to smooth the input image before segmenting it. (default: 0.5)

--content-threshold CONTENT_THRESHOLD

Threshold parameter indicating the proportion of the tile area that should be foreground (tissue content) in order to be selected. It should range between 0 and 1. (default: 0.5)

### Supplementary Figures

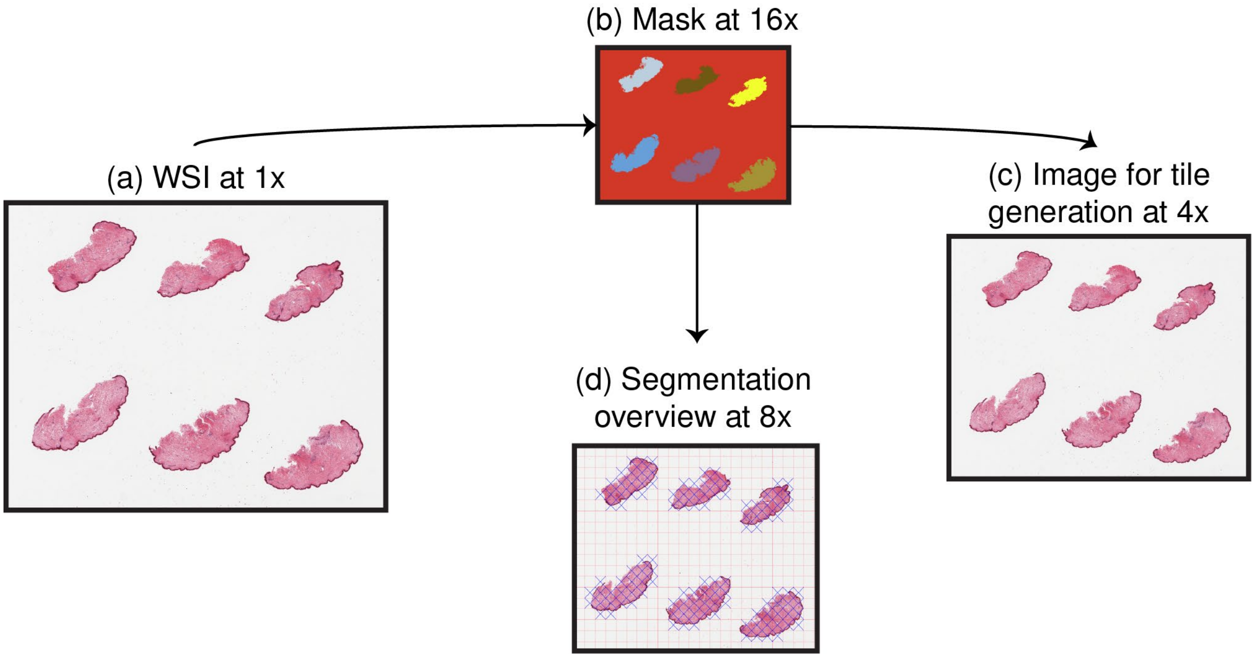


**Fig. S1: WSI scaling steps in PyHIST. (a)** WSI at its original resolution (1x). **(b)** The mask can be generated and processed at a given downsampling factor. A smaller resolution will lead to a faster segmentation. **(c)** The output can be requested at a given downsampling factor. **(d)** The segmentation overview image can also be generated at a given downsampling factor. The dimensions in all steps are matched to ensure that the tile sizes and grid are consistent. The downsampling choices for all the steps are independent of each other.

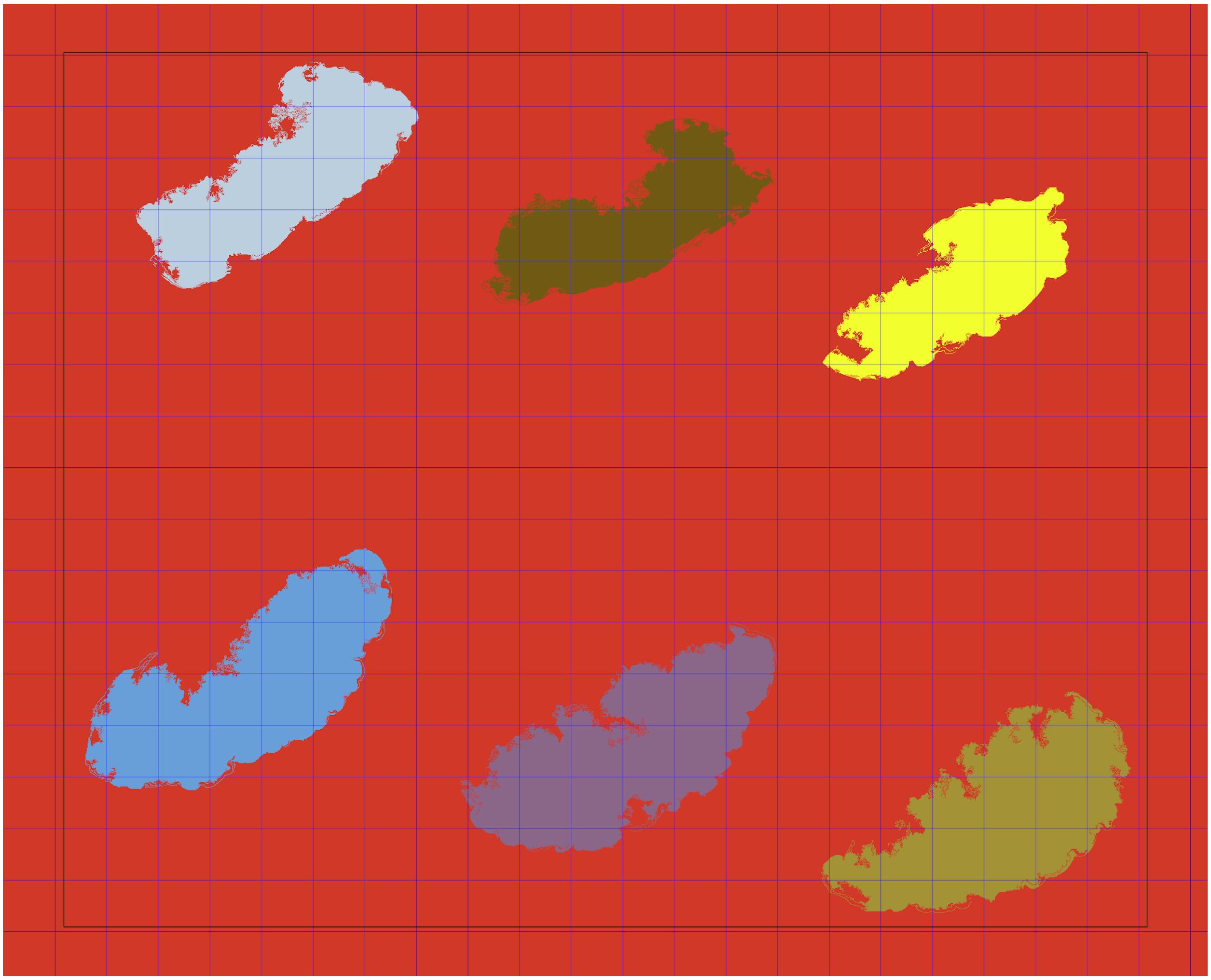

**Fig. S2: Image in test mode.** Test mode allows the user to see how the image mask will be with the chosen segmentation parameters and tile dimension configuration, before proceeding to generate the individual tile files. The black border defines the region of exclusion for tissue content placed within the edges of the slide (see *--borders* and *--corners* arguments).

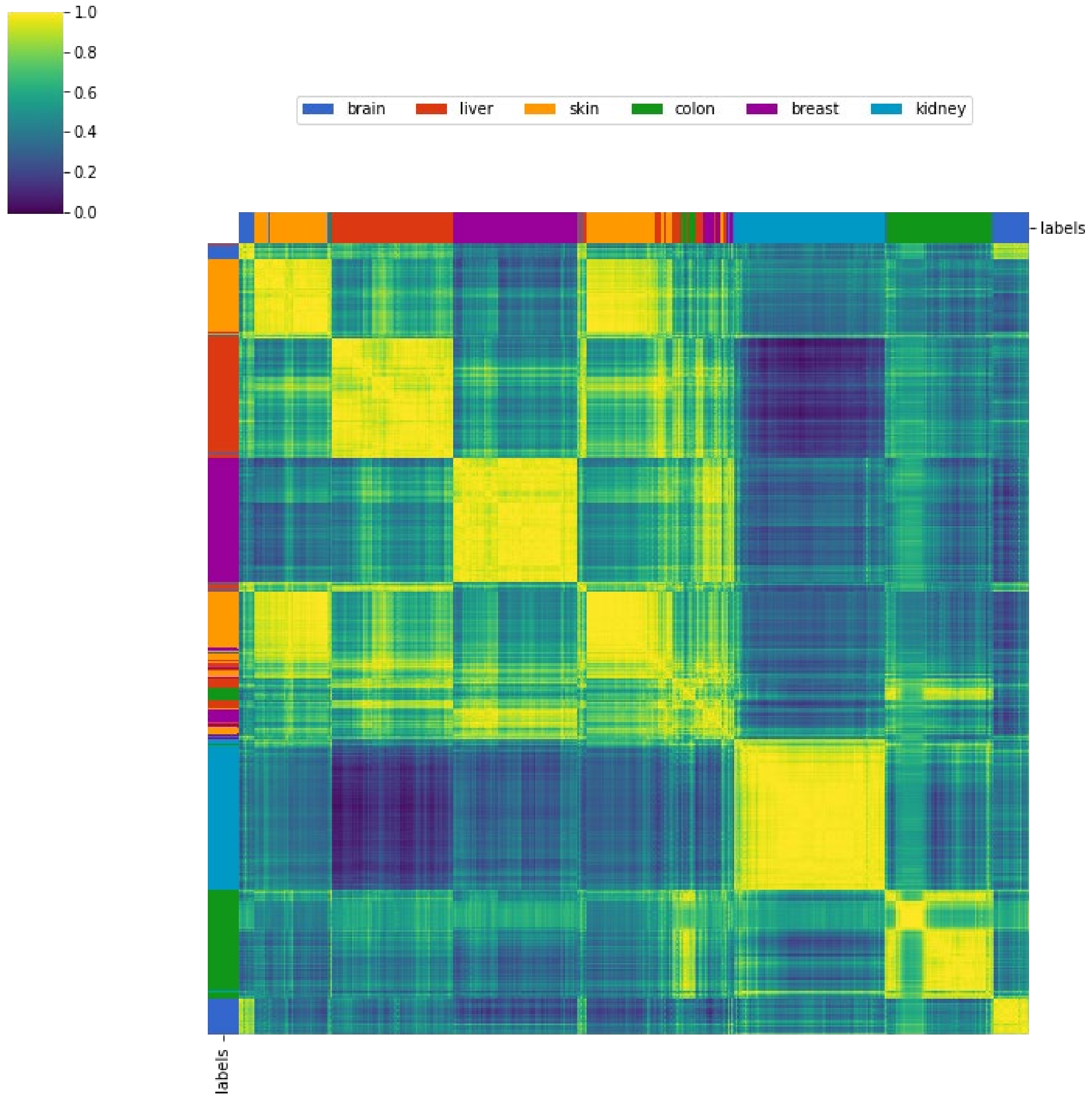


**Fig. S3: Correlation matrix of TCGA tiles based on their feature vectors.** Heatmap of Pearson's correlation matrix between the feature vectors obtained for each TCGA tile. Rows and columns are reordered with hierarchical agglomerative clustering.

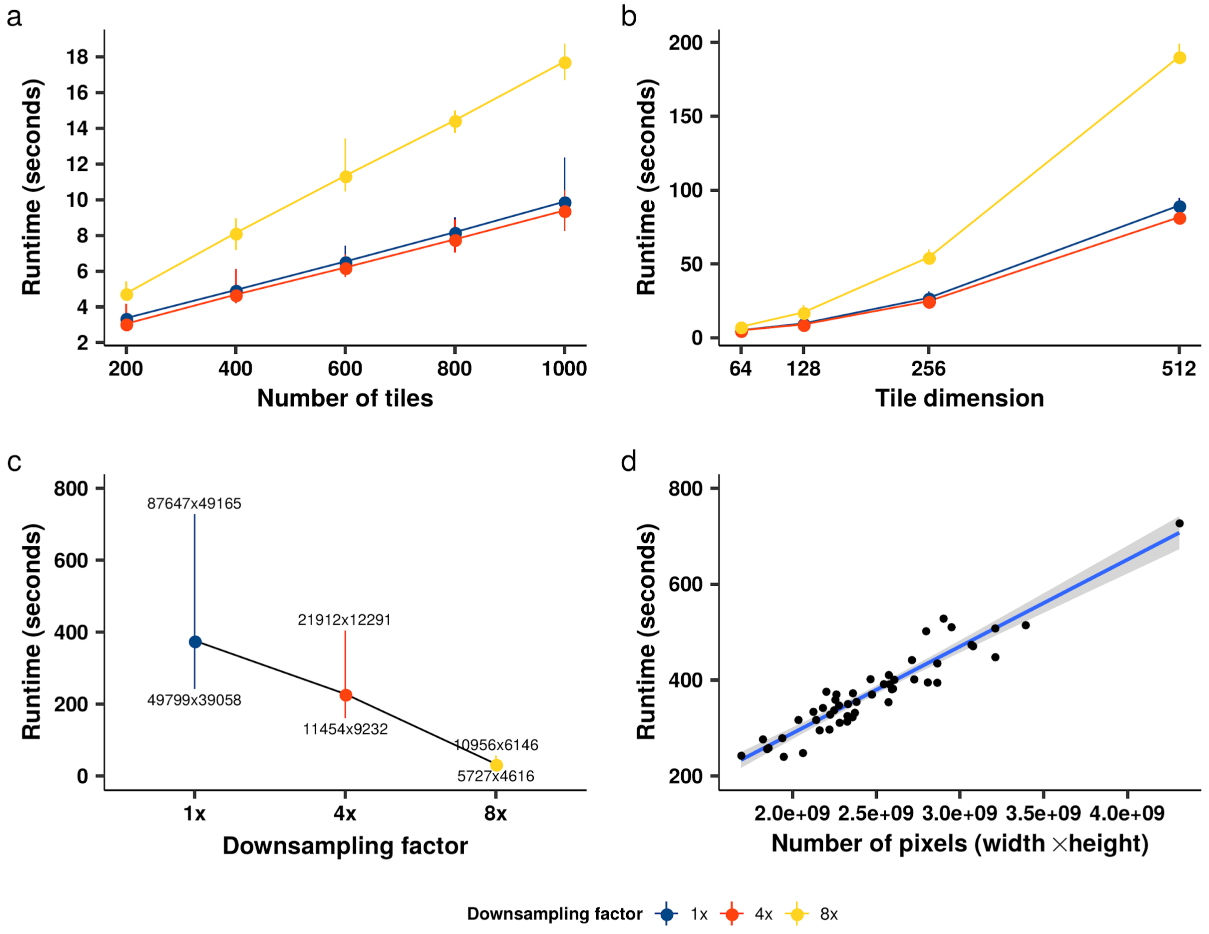


**Fig. S4: Runtime benchmarks. (a)** Execution time to perform random sampling (y-axis) of a varying number of tiles (x-axis) at different downsampling factors for the WSI shown in Fig. S1. For each combination of number of tiles and downsampling factor, the sampling was repeated 30 times. Each dot represents the average running time across the 30 runs, while the interval shows the range between the maximal and minimal running time. **(b)** Execution time to perform random sampling of 1000 tiles (y-axis) at different tile dimensions (x-axis) at different downsampling factors for the same WSI in (a). Each combination was repeated 50 times, with each dot showing the average runtime. **(c)** Segmentation runtime of 50 Stomach WSIs from the GTEx project, at different downsampling factors, at a tile size of 256x256. Each dot represents the average execution time. Each interval shows the range between the fastest and slowest segmentations, while the labels show the dimensions of the corresponding WSIs. **(d)** Segmentation runtime (y-axis) at 1x resolution for the 50 Stomach WSIs, with respect to the number of pixels in the WSI (x-axis).

### Supplementary Tables

| | | Predicted | | | | | |
|---|---|---|---|---|---|---|---|
| | | Brain | Breast | Colon | Kidney | Liver | Skin |
| **Real** | Brain | 94 | 2 | 3 | 0 | 0 | 11 |
| | Breast | 0 | 259 | 1 | 0 | 0 | 7 |
| | Colon | 0 | 0 | 205 | 0 | 0 | 6 |
| | Kidney | 1 | 0 | 0 | 271 | 0 | 0 |
| | Liver | 2 | 5 | 2 | 0 | 242 | 28 |
| | Skin | 1 | 3 | 1 | 1 | 2 | 269 |

**Table S1:** Confusion matrix for the tiles in the test set of the TCGA use case.
